# Investigating neurotrophin genetics and hippocampal volume

**DOI:** 10.1101/2022.01.01.474700

**Authors:** Jaisalmer de Frutos Lucas, Michael Vacher, Tenielle Porter, Belinda Brown, Simon Laws

## Abstract

As individuals get older, the structural integrity of brain regions becomes progressively diminished. Neurotrophic function might aid in preventing such losses through increased synaptogenesis and neurogenesis, particularly in the hippocampus, a brain structure relevant for cognitive function. However, the carriage of certain genetic alleles for genes involved in neurotrophic function might restrain the effectiveness of neurotrophin signalling, hindering neuroprotection. Yet, research on the contribution of single nucleotide polymorphisms (SNPs) within genes coding for neurotrophins and their receptors to hippocampal volumes is scarce, with the exception of rs6265 within the brain-derived neurotrophic factor gene. Therefore, the aim of this study was to identify SNPs within genes involved neurotrophic function that are associated with hippocampal volume in a sample of 23,776 cognitively normal older adults from the UK Biobank. We found that, in individuals older than 50, homozygote carriage of the major alleles rs4839435-A within nerve growth factor gene and rs56405676-T within the neurotrophic receptor tyrosine kinase 2 gene, were associated with increase hippocampal volumes, compared to carriage of 1 or 2 copies of the minor alleles. However, only rs56405676-T was significantly associated with greater hippocampal volumes in individuals older than 60. Hence this study might serve to identify populations at higher risk of hippocampal attrition and cognitive decline.

## Introduction

Through the ageing process, the integrity of grey and white matter structures is compromised^1^. Certain regions, such as the hippocampus, are more susceptible to the noxious effects of time^2,3^. Relevantly, hippocampal attrition predicts cognitive decline and dementia^4,5^, although the relationship between hippocampal volume and cognitive performance in older healthy individuals remains to be fully elucidated^6^. Therefore, to prevent cognitive decline, it is crucial to identify populations at higher risk of grey matter reductions as well as strategies to prevent neurodegeneration.

Neurotrophins are a family of signalling molecules that promote neurogenesis and synaptic plasticity in the brain^7^. They signal through various tyrosine kinase receptors (TKR) and the low affinity receptor p75. The beneficial effects of brain-derived neurotrophic factor (BDNF) up-regulation on hippocampal short and long-term potentiation as well as on neurogenesis and hippocampal volume probably represent the most consolidated findings within this field^8–11^. Conversely, reductions in neurotrophin levels in the ageing brain are associated with neuronal loss and greater incidence of age-related neurodegenerative disorders^12^.

Most of the studies looking into the relationship between neurotrophins and brain structural integrity have focused on the effect of a specific single nucleotide polymorphism (SNP) involved in the formation and transcription of a particular neurotrophin, most frequently BDNF, with very few addressing other neurotrophins^13^. For example, a meta-analysis showed that individuals carrying the *BDNF* Met/Met gene variant present lower hippocampal volume, in comparison with those carrying the Val/Val variant^13,14^. However, to identify populations at greater risk of age-related grey and white matter attrition it is fundamental to take a more holistic approach and consider the combined effect of multiple SNPs involved in neurotrophin function. Therefore, in this study, we aim to identify SNPs within genes associated to neurotrophic function that are associated with hippocampal volume (left, right and mean hippocampal volume) in cognitively normal older adults from the UK Biobank.

## Methods

### Study participants

The UK Biobank (http://ukbiobank.ac.uk/) is a large prospective cohort study which started in 2006, collecting health data on over 500,000 participants aged 40-69 at baseline. Ethical approval was provided by the NHS National Research Ethics Committee (11/NW/0382). The analyses here presented were carried out as a part of the UK Biobank application 45567. All participants provided informed consent prior to participation.

### Brain Imaging

Volumetric MRI data were acquired using a Siemens Skyra 3T scanner (Siemens Healthcare, Erlangen, Germany), with a standard 32-channel RF receive head coil according to UK Biobank procedures previously reported^15^. Briefly, the 3D MPRAGE T1-weighted images underwent acquired pre-processing and analysis using FSL packages (version 5, FMRIB Software Library, Oxford, England). The image-processing pipeline developed by UK Biobank was applied to generate imaging-derived phenotypes^16^.

### Genotyping

Genotying for the UK Biobank was conducted by Affymetrix on purpose-designed arrays; ∼50,000 samples were analysed using the BiLEVE Axiom array and ∼450,000 on the Affymetric UK Biobank Axiom array. Quality control practices were performed as previously described^17^. For the purpose of this study, we focused on SNPs within genes associated with neurotrophic function: *BDNF*, nerve growth factor (*NGF*), RE1 silencing transcription factor (*REST*), nerve growth factor receptor (*NGFR*), neurotrophic receptor tyrosine kinase 1 (*NTRK1*) and neurotrophic receptor tyrosine kinase 2 (*NTRK2*).

### Statistical analyses

Analysis of demographic data included median/counts and IQR/percentages for age, *APOE* ε4 carriage and education. Hippocampal volume data was normalised using the “--pheno-quantile-normalize” command in Plink. All analyses of SNP associations used a *dominant* model for the minor allele where minor allele homozygotes and heterozygotes were groups and assessed against major allele homozygotes. Linear models in Plink were used to investigate associations between hippocampal volume and SNPs using the model outlined below. Finally, all analyses were corrected for the False Discovery Rate (FDR) using the Benjamini & Hochberg (1995)^18^ step-up FDR control.

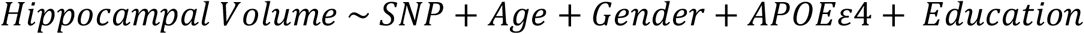

## Results

### Demographic information

23,776 UK Biobank cognitively normal participants over 50 years old with both genetic and MRI data were assessed in the current study. Demographic information for the UK Biobank sample utilised in the current study is presented in Table 1. A subset of these participants who were over 60 years old (n = 10,230) were also independently assessed (Table 1).

**Table 1.** >50-Year-Old Cohort Demographics

| | | >50 Years-Old<br>($n = 23,776$) | >60 Years-Old<br>($n = 10,230$) |
| --- | --- | --- | --- |
| Age |  | 58.0 (54.0, 62.0) | 63.0 (61.0, 65.0) |
| Sex | *Male* | 11,461 (48%) | 5,429 (53%) |
|  | *Female* | 12,315 (52%) | 4,801 (47%) |
| *APOE*<br>Genotype | *$\epsilon 4$ Negative* | 17,863 (75%) | 7,819 (76%) |
|  | *$\epsilon 4$ Positive* | 5,913 (25%) | 2,411 (24%) |
| Education | *College or University degree* | 10,877 (46%) | 4,281 (42%) |
|  | *A levels/AS levels or equivalent* | 2,918 (12%) | 1,073 (10%) |
|  | *O levels/GCSEs or equivalent* | 4,472 (19%) | 2,062 (20%) |
|  | *CSEs or equivalent* | 785 (3%) | 167 (2%) |
|  | *NVQ or HND or HNC or equivalent* | 1,353 (6%) | 688 (7%) |
|  | *Other professional qualifications*<br>*eg: nursing, teaching* | 1,349 (6%) | 732 (7%) |
|  | *None of the above* | 1,832 (8%) | 1133 (11%) |
|  | *Prefer not to answer* | 190 (1%) | 94 (1%) |

### SNPs associated with hippocampal volume

In the whole cohort (n = 23,776), 32 SNPs were nominally associated with left hippocampal volume, 31 with right hippocampal volume and 37 with mean hippocampal volume. Following correction for the false discovery rate rs4839435 within the *NGF* gene and rs56405676 within the *NTRK2* gene continued to be associated with mean, left and right hippocampal volume. For both SNPs homozygote carriage of the major alleles, rs4839435-A and rs56405676-T, were associated with increase hippocampal volumes when compared to those carrying 1 or 2 copies of the minor allele (Table 2).

**Table 2.** SNP Associations

| Cohort | Hippocampal<br>Volume<br>Measure | Chromosome | Position | rsID | Ref<br>Allele | Alt<br>Allele | $\beta$ | Nominal P | FDR P |
| --- | --- | --- | --- | --- | --- | --- | --- | --- | --- |
| >50 Years-Old |  |  |  |  |  |  |  |  |  |
|  | Mean | 1 | 115858104 | rs4839435 | G | A | 0.1126 | 5.46E-05 | 0.006 |
|  | Left | 1 | 115858104 | rs4839435 | G | A | 0.1090 | 0.0001 | 0.011 |
|  | Right | 1 | 115858104 | rs4839435 | G | A | 0.1064 | 0.0002 | 0.030 |
|  | Mean | 9 | 87573256 | rs56405676 | C | T | 0.0820 | 4.17E-05 | 0.006 |
|  | Left | 9 | 87573256 | rs56405676 | C | T | 0.0832 | 3.51E-05 | 0.008 |
|  | Right | 9 | 87573256 | rs56405676 | C | T | 0.0739 | 0.0003 | 0.030 |
| >60 Years-Old |  |  |  |  |  |  |  |  |  |
|  | Mean | 9 | 87573256 | rs56405676 | C | T | 0.0162 | 7.34E-05 | 0.123 |
|  | Left | 9 | 87573256 | rs56405676 | C | T | 0.0193 | 8.72E-05 | 0.123 |

In the cohort restricted to only those over 60 years of age (n = 10,230), 24 SNPs were nominally associated with left hippocampal volume, 28 with right hippocampal volume and 23 with mean hippocampal volume. Following correction for the false discovery rate only rs56405676 continued to be associated with mean (p = 0.016) and left (p = 0.019) hippocampal volume. With homozygote carriage of the rs56405676 major allele (T) associated with increased hippocampal volumes when compared to those carrying 1 or 2 copies of the minor allele (C) (Table 2).

## Discussion

Neurotrophins are signalling molecules that play a major role in synaptogenesis and neurogenesis^7^. Current research supports the notion that the SNP *BDNF* rs6265 significantly contributes to individual differences in hippocampal volume^13,14,19^. Nevertheless, little is known regarding how genetic variance within other genes relevant to neurotrophic function might influence hippocampal structure and function. In this study, we provide novel evidence of the relevance of two SNPs associated with hippocampal volume in older adults: rs4839435 within the *NGF* gene and rs56405676 within the *NTRK2* gene. However, in the case of individuals older than 60, only rs56405676 within the *NTRK2* remained significantly associated with hippocampal volume. Surprisingly, no SNP within *BDNF* (nor any of the other genes explored) showed a significant association with hippocampal volume after FDR correction.

Previous research had identified several polymorphisms within the *NTRK2* gene associated with depression^20–22^. Interestingly, a significant relationship between hippocampal volume and depression had also been reported^23^. Yet, to the best of our knowledge, this is the first study to provide evidence of a direct association between a SNP within *NTRK2* and hippocampal volume. In fact, to the best of our knowledge no previous study has found a significant association between the specific SNP rs56405676 and any phenotypes. Regarding NGF, NGF deprivation had been described to induce basal forebrain and hippocampal atrophy, which had been associated to cognitive decline and Alzheimer’s disease^24^. In this line, a different SNP within *NGF*, rs6330, had been linked to poorer executive function in early-stage Alzheimer’s disease and mild cognitive impairment^25^. However, as in the case of rs56405676 within *NTRK2*, rs4839435 within *NGF* had not been previously associated with hippocampal structural or functional traits, or any other phenotypes. Whether these newly identified SNPs impair the expression and function of NGF and NTRK2 in the hippocampus remains to be investigated. If that was the case, future studies should consider the clinical implications of such alterations. Moreover, several factors (i.e., environmental enrichment, physical activity) have been suggested to up-regulate the expression of neurotrophins^26–28^. Future studies could also address whether greater engagement in such activities compensates for the detrimental effects of rs4839435-A and rs56405676-T carriage on hippocampal volume or whether the carriage of those genetic variants hinders the lifestyle-induced up-regulation of NGF and NTRK2.

## Notes

### Competing Interest Statement

The authors have declared no competing interest.

### Summary of Updates

List of authors

